# Linear ubiquitination triggers Amph-mediated T-tubule biogenesis

**DOI:** 10.1101/2025.04.03.646909

**Authors:** Kohei Kawaguchi, Yutaro Hama, Harunori Yoshikawa, Kohei Nishino, Kazuki Morimoto, Tsuyoshi Nakamura, Michiko Koizumi, Yuriko Sakamaki, Kota Abe, Soichiro Kakuta, Koichiro Ichimura, Fumiyo Ikeda, Hidetaka Kosako, Naonobu Fujita

## Abstract

T-tubules are specialized invaginations of the plasma membrane essential for muscle contraction. While their physiological importance is well established, the mechanisms underlying T-tubule formation remain elusive. Here, we identify LUBEL/RNF31, a ubiquitin E3 ligase responsible for linear (M1-linked) ubiquitination, as a key regulator of T-tubule biogenesis through proximity proteomics and RNAi screening in *Drosophila*. Loss of LUBEL leads to the formation of Amphiphysin (Amph)-positive membrane sheets instead of tubular networks in muscle cells. Mechanistically, the ubiquitin ligase activity of LUBEL, and direct interaction with Amph, a BAR domain protein involved in membrane tubule extension, are crucial for proper T-tubule morphology. LUBEL and M1-linked ubiquitin chains assemble into condensates on membranes, facilitating Amph-mediated membrane tubulation. Notably, the Amph-LUBEL/RNF31 interaction is evolutionarily conserved across a broad range of species, underscoring a fundamental role for linear ubiquitination in membrane remodeling. Our findings uncover a previously unrecognized role for linear ubiquitination in membrane deformation driven by BAR domain proteins.

## Introduction

Cellular membrane function requires continuous remodeling orchestrated by the coordinated actions of lipids and curvature-generating proteins, such as caveolins and BAR domain-containing proteins (*1–4*). Recent studies have demonstrated that protein condensates, formed through multivalent interactions, can drive membrane deformation via wetting, molecular crowding, and viscoelastic effects (*5–8*). These emerging concepts add a new layer of complexity to our understanding of the molecular mechanisms shaping membranes.

Protein condensation often relies on intrinsically disordered regions (IDRs), which mediate weak intra- and intermolecular interactions, and protein-interacting motifs that contribute to multivalency (*9*). A key regulatory mechanism of protein condensation is ubiquitination, a post-translational modification in which ubiquitin is covalently attached to a substrate (*10*, *11*). Ubiquitination can alter the molecular configuration of target proteins and provide additional binding valency for partners containing ubiquitin-binding domains (*12*). Given that ubiquitination is reversible, it may dynamically regulate membrane remodeling through condensation in a spatiotemporal manner. However, the interplay between ubiquitin-containing condensates and curvature-generating proteins remains largely unexplored.

Transverse tubules (T-tubules) are specialized plasma membrane invaginations that form a unique tubular network in muscle cells (*13*). During excitation-contraction coupling, a stimulus from motor neurons triggers depolarization of sarcolemma, the plasma membrane in muscle cells. This depolarization propagates through the T-tubules, ensuring the rapid and synchronized transmission of signals to the sarcoplasmic reticulum (SR), which releases calcium ions into the cytosol, ultimately driving muscle contraction and movement. Defects in T-tubule organization are linked to various congenital muscular diseases, underscoring their physiological significance.

Many key regulators of T-tubule shaping have been identified as causative genes of congenital myopathies (*14*). Centronuclear myopathy (CNM), a severe congenital myopathy characterized by T-tubule defects and muscle weakness, is associated with mutations in three membrane trafficking-related genes: BIN1 (Bridging Integrator-1; also known as Amph2 (Amphiphysin 2)), DNM2 (Dynamin 2) and MTM1 (Myotubularin 1) (*14*). Amph2 is a BAR domain protein that induces membrane tubulation (*15–18*). Its N-terminal BAR domain forms banana-shaped dimers that interact with negatively charged phospholipids via their positively charged concave surfaces. These dimers further assemble into extensive networks, promoting membrane curvature and tubulation (*19*). The C-terminal SH3 domain of Amph2 acts as an adapter, binding to partner proteins such as DNM2, which is a large GTPase involved in membrane scission (*20*). Excessive DNM2 activity disrupts the T-tubule network (*21*), indicating the necessity of precisely regulated DNM2 activity for proper T-tubule formation. The maintenance of the T-tubule network involves MTM1, a phosphatidylinositol 3-phosphate (PI3P) phosphatase (*22*). Additionally, the muscle-specific caveolae-forming protein Cav3 has been implicated in the early stage of T-tubule formation and is associated with several myopathies, including limb-girdle muscular dystrophy (LGMD1C). The caveolin-associating coat protein, Cavin4, is also required for T-tubule formation and has been identified as a causative gene for dilated cardiomyopathy (*23*, *24*). Moreover, its paralog, Cavin1, has been shown to contribute to T-tubule formation in mice and zebrafish (*25*). Cavin4 has recently been reported to bridge caveolae and Amph2, forming a ring-like platform at the sarcolemma that acts as a scaffold for initiation of T-tubule formation (*23*, *24*). However, while the caveola-forming caveolins Cav1 and Cav3 are conserved in chordates and vertebrates, they are absent in other metazoans (*26*). Remarkably, T-tubules emerged evolutionarily before the caveolin-mediated membrane deformation mechanisms (*27–29*), suggesting the existence of an alternative: a caveolin-independent mechanism for T-tubule biogenesis.

*Drosophila* has emerged as an ideal model organism for studying T-tubule biogenesis. Several characteristics make *Drosophila* particularly suited for T-tubule research: first, T-tubules are not essential for viability in flies (*17*); second, T-tubules can be observed through the cuticle in live animals; and third, *Drosophila* is genetically tractable. These advantages enable reverse genetic analysis, which is essential for elucidating mechanisms of T-tubule formation. This study employed proximity labeling, an enzymatic reaction that enables the identification of local proteomes, and *in vivo* RNAi screening to identify key T-tubule regulators. Our analysis revealed that LUBEL, an E3 ligase responsible for linear (M1-linked) ubiquitination, plays a crucial role in T-tubule biogenesis. LUBEL promotes the formation of M1-linked ubiquitin chains, in which the α-amino group of the N-terminal methionine (M1) of ubiquitin is used instead of the ε-amino group of lysine residues (K6, K11, K27, K29, K33, K48, and K63) (*12*, *30*, *31*). Further mechanistic investigations into LUBEL uncovered an unexpected role of linear ubiquitination in BAR protein-mediated membrane remodeling.

## Results

### Proximity proteomics of T-tubules

We previously established an imaging-based RNAi screening system for T-tubules in *Drosophila* larval body wall muscles (BWMs), which possess developed T-tubules oriented both longitudinally and transversely (Fig. 1A) (*32*). While genome-wide RNAi screening is feasible in *Drosophila*, it is time- and labor-intensive. To identify candidate genes unbiasedly, we employed biotinylation-based proximity proteomics of T-tubules *in vivo*. We generated a transgenic line carrying an Amph construct fused with miniTurbo (mnTb), an engineered biotin ligase designed for proximity labeling (*33*). Immunostaining confirmed that Amph-mnTb, but not mnTb alone, colocalized with Dlg1, an established T-tubule marker, in larval muscle cells (Fig. S1A), in the same manner as endogenous Amph (*17*, *32*).

**Figure 1.**
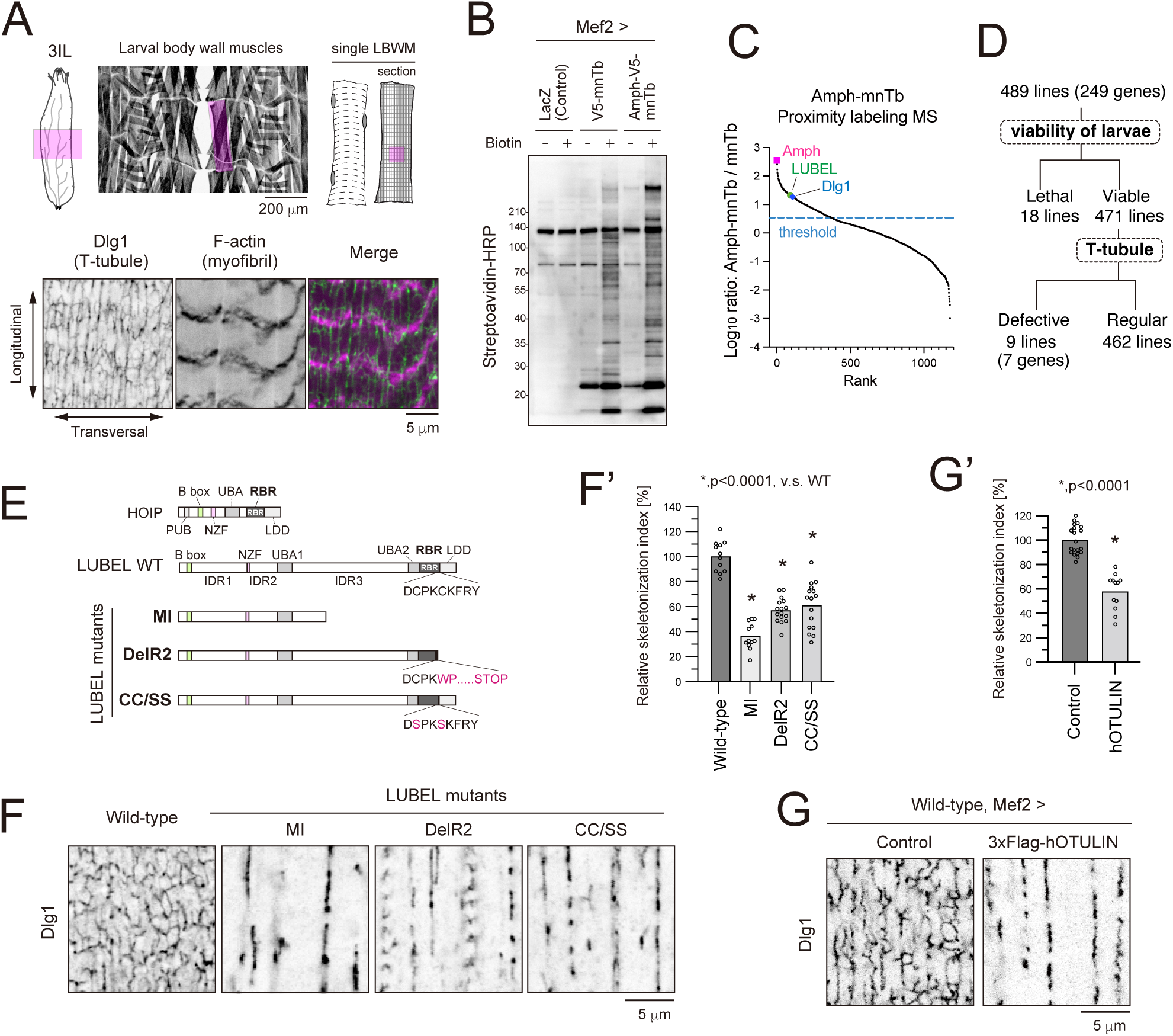
Proximity proteomics and RNAi screening identify LUBEL as a critical regulator of T-tubule formation. (A) T-tubules in the third-instar larva (3IL) body wall muscle (BWM). Representative images of T-tubules (Dlg1, green) and myofibrils (F-actin, magenta) in the midsection of 3IL BWM. (B) Streptavidin blotting of larval carcass fillet lysates. LacZ, mnTb, or Amph-mnTb were expressed in BWMs. Larvae were reared with or without biotin for 2 days before sample preparation. (C) Proximity proteomics of T-tubules using Amph-mnTb. The Y-axis represents the fold change (Amph-mnTb vs. mnTb control), while the X-axis indicates rank. (D) Summary of RNAi screening results. (E) Schematic representation of LUBEL mutants. LUBEL contains the indicated domains. LUBEL mutants either lack the C-terminal region or harbor specific mutations. (F, F’) T-tubules in 3IL BWMs of indicated genotypes. Images of anti-Dlg1 staining (F) and quantification of skeletonized Dlg1-positive structures (F’). (G, G’) Effect of human OTULIN overexpression on T-tubules. Images of anti-Dlg1 staining (G) and quantification of skeletonized Dlg1-positive structures (G’).

To verify the biotinylation capability of Amph-mnTb, we expressed either Amph-mnTb or mnTb alone in muscles, and larvae were reared in the presence or absence of biotin. As a control, LacZ was expressed instead of the mnTb-fused constructs. Streptavidin-HRP blotting of larval carcass lysates revealed intense biotin-dependent labeling in Amph-mnTb and mnTb samples but not in controls (LacZ) (Fig. 1B). Notably, distinct banding patterns between Amph-mnTb and mnTb samples were observed under biotin-feeding conditions, suggesting that Amph-mnTb biotinylates its proximal proteins. Biotinylated peptides were purified from trypsin-digested samples using Tamavidin2-REV beads and analyzed by mass spectrometry (*34*), revealing the proximity proteome of Amph in muscle cells (Data S1). As expected, Dlg1 was significantly enriched in Amph-mnTb samples (Fig. 1C and Data S1), confirming successful T-tubule proximity labeling.

### Muscle-targeted RNAi screening for T-tubule regulators

From the proximity proteomics data (Data S1), we selected 249 genes surpassing the enrichment threshold for muscle-targeted RNAi screening for T-tubule regulators (Fig. 1D). A total of 489 UAS-RNAi lines were crossed with a line carrying both Dlg1-GFP, a T-tubule marker driven by its endogenous promoter, and Mef2-GAL4. Of these, 18 RNAi lines resulted in lethality, thus the remaining 471 viable lines were further analyzed. Confocal microscopy imaging of Dlg1-GFP-labeled T-tubules in intact larvae identified nine RNAi lines exhibiting defects in T-tubule morphology (Fig. 1D and Data S2).

Among them, three independent RNAi lines targeting Linear Ubiquitin E3 ligase (LUBEL) induced the most severe phenotype, resembling the T-tubule defects observed in Amph RNAi (Fig. S1B). Both conditions resulted in the loss of transverse and thick longitudinal membranes labeled with Dlg1. LUBEL is the *Drosophila* ortholog of HOIP, the catalytic subunit of the linear ubiquitin chain assembly complex (LUBAC) in mammals (*30*). LUBAC is composed of three subunits: HOIP (HOIL-1-interacting protein), HOIL-1 (heme-oxidized IRP2 ubiquitin ligase 1), and SHARPIN (SHANK-associated RH domain-interacting protein). Among these, HOIP acts as the catalytic subunit responsible for M1-linked ubiquitination, while HOIL-1 and SHARPIN function as regulatory subunits that stabilize and regulate LUBAC activity (*35–39*). Unlike its mammalian counterpart, *Drosophila* LUBEL is sufficient for M1-linked ubiquitination without additional regulatory subunits (*30*). LUBEL shares multiple conserved domains with HOIP, including the B-box, Npl4 Zinc Finger (NZF), ubiquitin-associated (UBA), RING-between-RING (RBR), and linear ubiquitin determinant (LDD) domains (Fig. 1E). In addition, unlike HOIP, LUBEL contains extended IDRs between key domains (*30*), contributing to its larger size (2902 aa vs. 1072 aa in human HOIP). While M1-linked ubiquitination is critical for immune signaling pathways via NF-κB, its role in membrane remodeling remains unexplored. Given the pronounced T-tubule defects observed upon LUBEL knockdown (Fig. S1B), we focused on its function in subsequent analyses.

### M1-linked ubiquitination by LUBEL is indispensable for T-tubule morphology

To validate the RNAi-based knockdown phenotype, we analyzed a genomic mutant, *LUBEL^MI^*, which harbors a stop codon in the middle of the protein, leading to the loss of the entire C-terminal region, including the RBR domain (Fig. 1E). This mutant exhibited severe T-tubule defects (Fig. 1F and F’) confirming that LUBEL is essential for T-tubule formation. Next, we assessed whether the E3 ligase activity of LUBEL was required for T-tubule formation by analyzing mutant strains carrying enzymatically inactive variants (Fig. 1E). The *LUBEL^DelR2^* mutant, with a truncation of the C-terminal region of the RBR, and the *LUBEL^CC/SS^*mutant, carrying mutations in essential cysteine residues, both displayed abnormal T-tubule morphology (Fig. 1F), demonstrating that the E3 ligase activity of LUBEL is critical for T-tubule formation.

To determine whether M1-linked ubiquitination, rather than other ubiquitin linkages, is essential for T-tubule formation, we overexpressed human OTULIN, a deubiquitinating enzyme specific for M1-linked ubiquitin chains (*40*), in wild-type *Drosophila* muscle cells. Notably, OTULIN overexpression recapitulated the T-tubule defects observed in LUBEL loss-of-function mutants (Fig. 1G and G’), indicating an essential role of M1-linked ubiquitin chains in T-tubule formation. Together, these findings demonstrate that LUBEL-mediated linear ubiquitination is crucial for T-tubule formation.

### T-tubule formation is independent of the NF-κB signaling pathway

LUBAC-mediated M1-linked ubiquitination is well-established as a key regulator of the NF-κB signaling pathway in vertebrates (*31*, *41*). LUBAC ubiquitinates NEMO (NF-κB essential modulator), a regulatory subunit of the IKK (IκB kinase) complex, leading to the activation of NF-κB transcription factors (*35–37*, *39*). The role of M1-liked ubiquitination in NF-κB signaling is also evolutionarily conserved in *Drosophila* (*42*). To investigate if the NF-κB signaling pathway is required for T-tubule formation, we analyzed T-tubule morphology in loss-of-function mutants of this pathway. *Rel* encodes the *Drosophila* NF-κB protein, and *Kenny* is the *Drosophila* ortholog of IKKγ/NEMO (*43*, *44*). Both loss-of-function mutants, *Rel^E20^*and *Kenny^c02831^*, exhibited normal T-tubule morphology (Fig. S2), suggesting that NF-κB signaling is dispensable for T-tubule formation.

### LUBEL plays a critical role in membrane tubule extension mediated by Amph

Amph shapes tubular morphology through its membrane curvature activity (*15–18*). Based on the evidence that loss of LUBEL phenocopies Amph RNAi (Fig. S1B), we hypothesized that loss of LUBEL leads to a decrease in Amph protein levels, resulting in T-tubule defects. To test this, we performed immunoblotting of larval carcass lysates using anti-Amph antibodies (Fig. 2A). In the control sample, three Amph splicing isoforms―A, B, and C―were detected.

**Figure 2.**
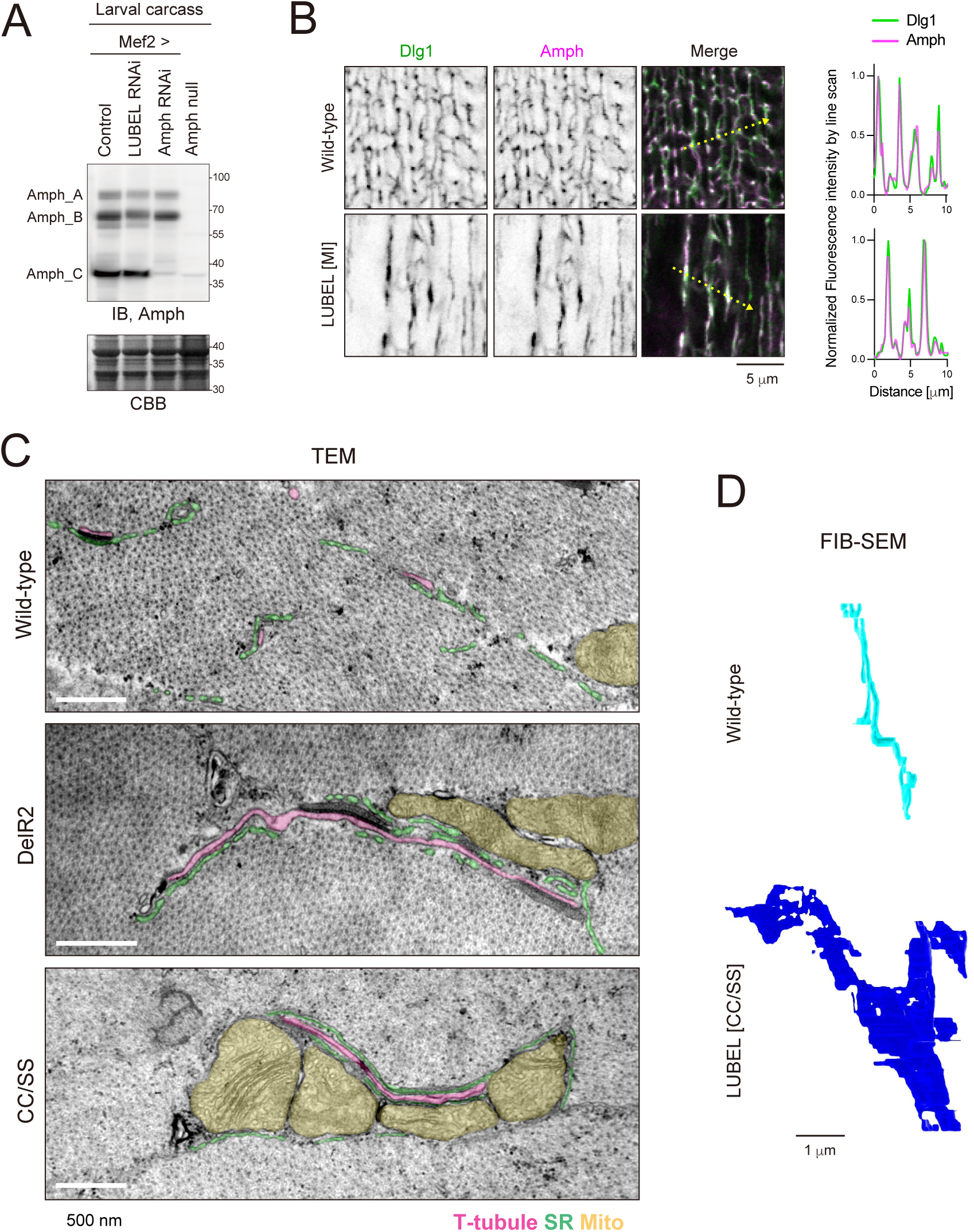
LUBEL is essential for Amph-dependent membrane tubulation. (A) Amph immunoblot of 3IL carcass fillet lysates from the indicated genotypes. Coomassie Brilliant Blue (CBB) staining was used as a loading control. (B) Immunostaining of Amph (magenta) and Dlg1(green) in wild-type and LUBEL mutant 3IL BWMs. Line plot profiles along the yellow dotted arrow are shown. (C) TEM images of wild-type and LUBEL mutant 3IL BWMs. T-tubules (magenta), sarcoplasmic reticulum (SR, green), and mitochondria (yellow) are highlighted. (D) 3D reconstruction models generated by FIB-SEM. T-tubules in wild-type (cyan) and T-tubule-related membranes in LUBEL mutant (blue) are shown.

Muscle-targeted Amph knockdown using Mef2-GAL4 specifically reduced Amph_C levels without affecting isoforms A or B, indicating that Amph_C is predominantly expressed in larval muscle cells. Contrary to our assumption, LUBEL knockdown did not alter Amph_C protein levels (Fig. 2A). These results indicate that the T-tubule defect caused by LUBEL deficiency is not due to changes in Amph protein levels.

Next, we investigated whether LUBEL influences Amph membrane localization by performing immunostaining for endogenous Amph in wild-type and LUBEL mutant muscle cells. As previously reported, Amph colocalized with Dlg1 on T-tubules in wild-type cells (Fig. 2B) (*17*, *32*). And while we observed altered Dlg1-positive membrane morphology, Amph and Dlg1 remained colocalized in LUBEL mutant muscle cells (Fig. 2B). This suggests that LUBEL does not affect the membrane localization of Amph but may regulate its membrane-bending function.

Although confocal microscopy revealed defects in Dlg1-positive membranes in LUBEL mutants, their ultrastructure remained unclear. To explore their ultrastructure, we performed transmission electron microscopy (TEM) imaging of cross-sections of larval muscle cells from wild-type and LUBEL mutants (Fig. 2C). In wild-type muscle cells, circular T-tubule cross-sections (highlighted in pink) were frequently observed. In contrast, LUBEL mutants exhibited elongated, tube-like structures (pink), which may have originated from T-tubules or their precursors. Additionally, these tubular structures were often flanked by the SR (green) and mitochondria (yellow), suggesting the importance of T-tubules in the spatial organization of the SR and mitochondria within muscle cells.

To further elucidate the three-dimensional (3D) architecture of T-tubule-related membranes in LUBEL mutants, we conducted focused ion beam scanning electron microscopy (FIB-SEM). T-tubules and their derived membranes were segmented in individual SEM images and reconstructed into 3D models. In wild-type muscle cells, T-tubules exhibited a tubular morphology, as expected. In contrast, T-tubule-related membranes in LUBEL mutants appeared as flat, sheet-like structures (Fig. 2D, Movies S1 and S2). These findings collectively indicate that LUBEL promotes membrane tubule extension without affecting Amph protein levels.

### LUBEL directly interacts with Amph to form T-tubules

LUBEL contains multiple domains besides the RBR domain (Fig. 3A). To assess whether its N-terminal region contributes to T-tubule biogenesis, we performed rescue experiments using either a full-length LUBEL construct (Full) or one lacking the N-terminal region (LUBEL^873-2902^) (Fig. 3A). The full-length construct rescued the T-tubule defects in the *LUBEL^DelR2^* mutant, whereas LUBEL^873-2902^ did not (Fig. 3B and B’). Moreover, overexpression of the N-terminal fragment (LUBEL^1-872^) disrupted the T-tubule morphology even in wild-type muscle cells (Fig. 3C and C’). Furthermore, LUBEL^1-872^ localized to Dlg1-positive membranes (Fig. S3A), suggesting that the N-terminal region may sequester an unknown factor required for membrane localization. To identify potential binding partners, we performed immunoprecipitation-mass spectrometry (IP-MS) of LUBEL^1-872^ (Fig. 3D and Data S3) and proximity proteomics using mnTb-fused LUBEL (Fig. 3E and Data S4). Both analyses consistently identified Amph as a high-confidence candidate interactor of LUBEL.

**Figure 3.**
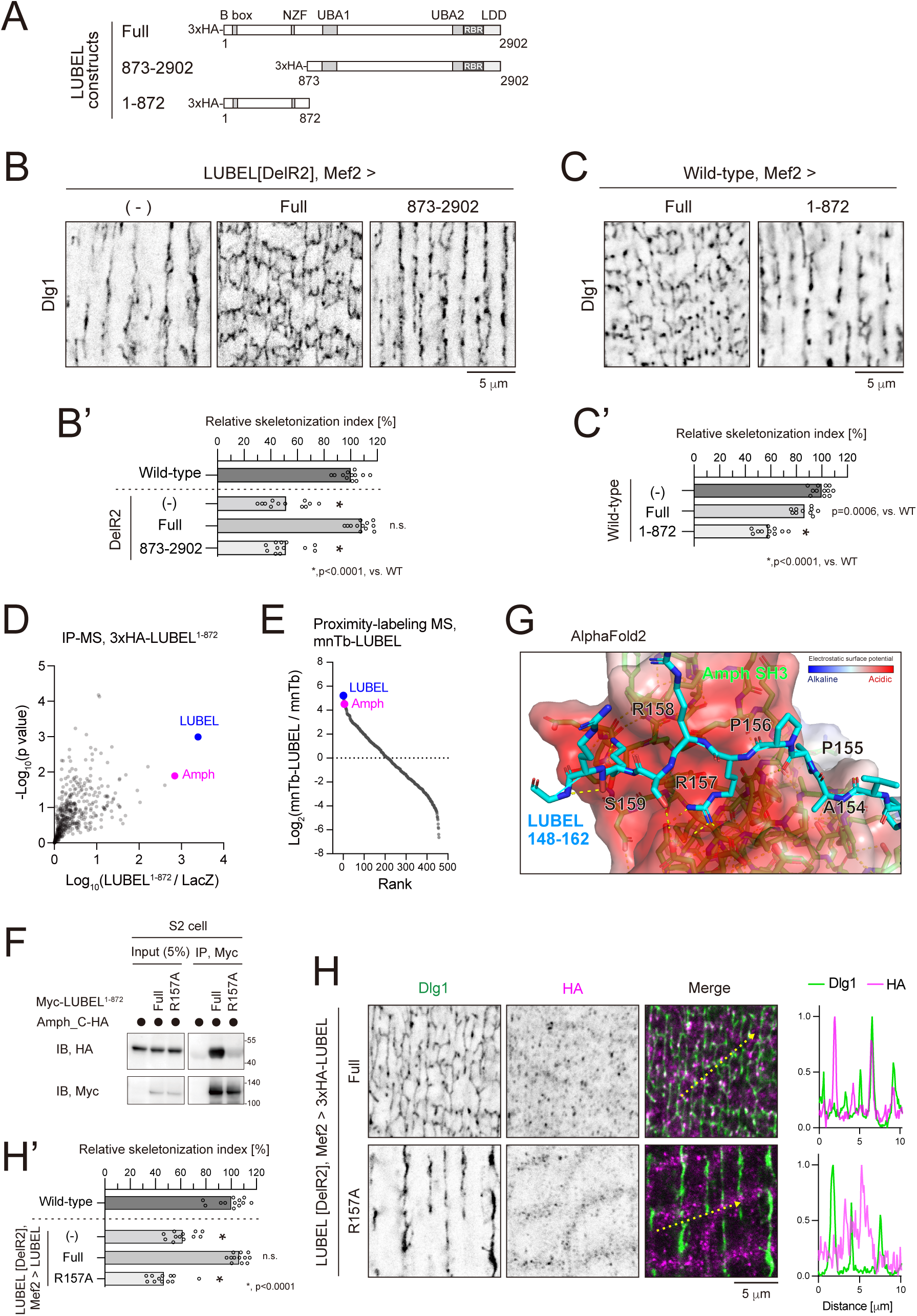
LUBEL-Amph interaction is essential for T-tubule formation. (A) Schematic representation of LUBEL truncations. (B, B’) LUBEL rescue experiment with full length protein (Full) or a construct lacking the N-terminal region (873–2902). 3IL BWMs were stained for Dlg1 (B), and their relative skeletonization index was quantified (B’). (C, C’) Effect of the LUBEL N-terminal fragment overexpression on T-tubules. Full-length protein (Full) or its N-terminal fragment (1–872) were expressed in wild-type 3IL BWM followed by staining for Dlg1 (C). HA immunostaining images are shown in Fig. S3A. The relative skeletonization index is shown (C’). (D) IP-MS analysis using HA-LUBEL^1-872^. HA-LUBEL^1-872^ or LacZ (control) was expressed in 3IL BWMs. Lysates from carcass fillets were subjected to anti-HA IP and analyzed by LC-MS/MS. The X-axis shows the fold change (HA-LUBEL^1-872^ vs. LacZ control), while the Y-axis represents the negative log10 P-value from two replicates. (E) Proximity proteomics using mnTb-LUBEL. The Y-axis represents the fold change (mnTb-LUBEL vs. mnTb control), while the X-axis indicates rank. (F) Co-IP assay of Amph and LUBEL^R157A^. Myc-LUBEL^1-872^ (wild-type or R157A mutant) and Amph isoform C-HA were co-expressed in S2 cells. Lysates were subjected to anti-Myc IP and immunoblotted for anti-Myc and anti-HA antibodies. (G) Structure of the Amph SH3 and LUBEL-N fragment complex using AlphaFold 2. For LUBEL, only residues 148–168 are displayed. (H, H’) LUBEL rescue experiments using full-length protein (Full) or the LUBEL^R157A^ (R157A) mutant. 3IL BWMs were stained with anti-Dlg1 and anti-HA antibodies (H). The relative skeletonization index is shown (H’).

To map the LUBEL-binding region in Amph, we performed GST pulldown assays using truncated Amph constructs (Fig. S3B). The C-terminal SH3 domain, but not the N-terminal BAR domain, pulled down LUBEL^1-872^ (Fig. S3C), indicating that the SH3 domain mediates the interaction between LUBEL and Amph. *In silico* analysis using AlphaFold2 predicted that four proline-rich motifs and the B-box zinc finger domain in LUBEL^1-872^ are binding sites for the Amph SH3 domain (*45*) (Fig. S3D). To validate this, we generated LUBEL point mutants designed to disrupt these interactions (Fig. S3E) and performed co-IP assays with Amph (Fig. S3E’). The LUBEL R157A mutation, an alanine substitution of Arg157, abolished binding with Amph (Fig. 3F and S3E’), identifying the proline-rich motif (152–157) as the binding site for the Amph SH3 domain. AlphaFold2 multimer modeling predicted that LUBEL Arg157 is buried in a 139.8 Å^2^ surface area of the Amph SH3 domain, which is 29.6% of the total binding interface (472.0 Å^2^). Arg157 is positioned to bind within an acidic pocket of the Amph SH3 domain with three Glutamate and Aspartate residues, and forms several salt bridges and hydrogen bonds with these acidic residues (Fig. 3G). These results suggest that the electrostatic interactions between the Arg157-containing proline-rich motif and the SH3 domain are important for LUBEL-Amph interaction.

To assess the functional significance of the LUBEL-Amph interaction in T-tubule formation, we expressed the LUBEL^R157A^ in LUBEL-deficient muscle cells. Unlike the full-length protein, LUBEL^R157A^ failed to rescue T-tubule defects (Fig. 3H and H’). Moreover, LUBEL R157A did not colocalize with Dlg1 (Fig. 3H). These findings highlight the critical role of the interaction between LUBEL and Amph in T-tubule formation.

### LUBEL and M1-linked ubiquitin chains form puncta on T-tubules

To examine the subcellular localization of LUBEL, we expressed mNeonGreen (mNG)-tagged LUBEL in muscle cells. mNG-LUBEL did not exhibit tubular network morphology but formed punctate structures that colocalized with Dlg1 (Fig. 4A). These LUBEL-positive puncta also contained M1-linked ubiquitin chains (Fig. 4B). Given its localization and role in membrane tubulation, LUBEL likely localizes to specific sites during T-tubule formation. However, direct observation of T-tubule formation is challenging due to its concurrence with myofibril differentiation. To overcome this issue, we utilized the GeneSwich GAL4 (GS) system to control mNG-LUBEL expression in LUBEL mutants temporally (*46*). Feeding larvae with RU486 (also known as mifepristone), an inducer of the GS system, triggers mNG-LUBEL expression. In the absence of RU486, Dlg1 appeared in thick, sheet-like structures (Fig. 4C).

**Figure 4.**
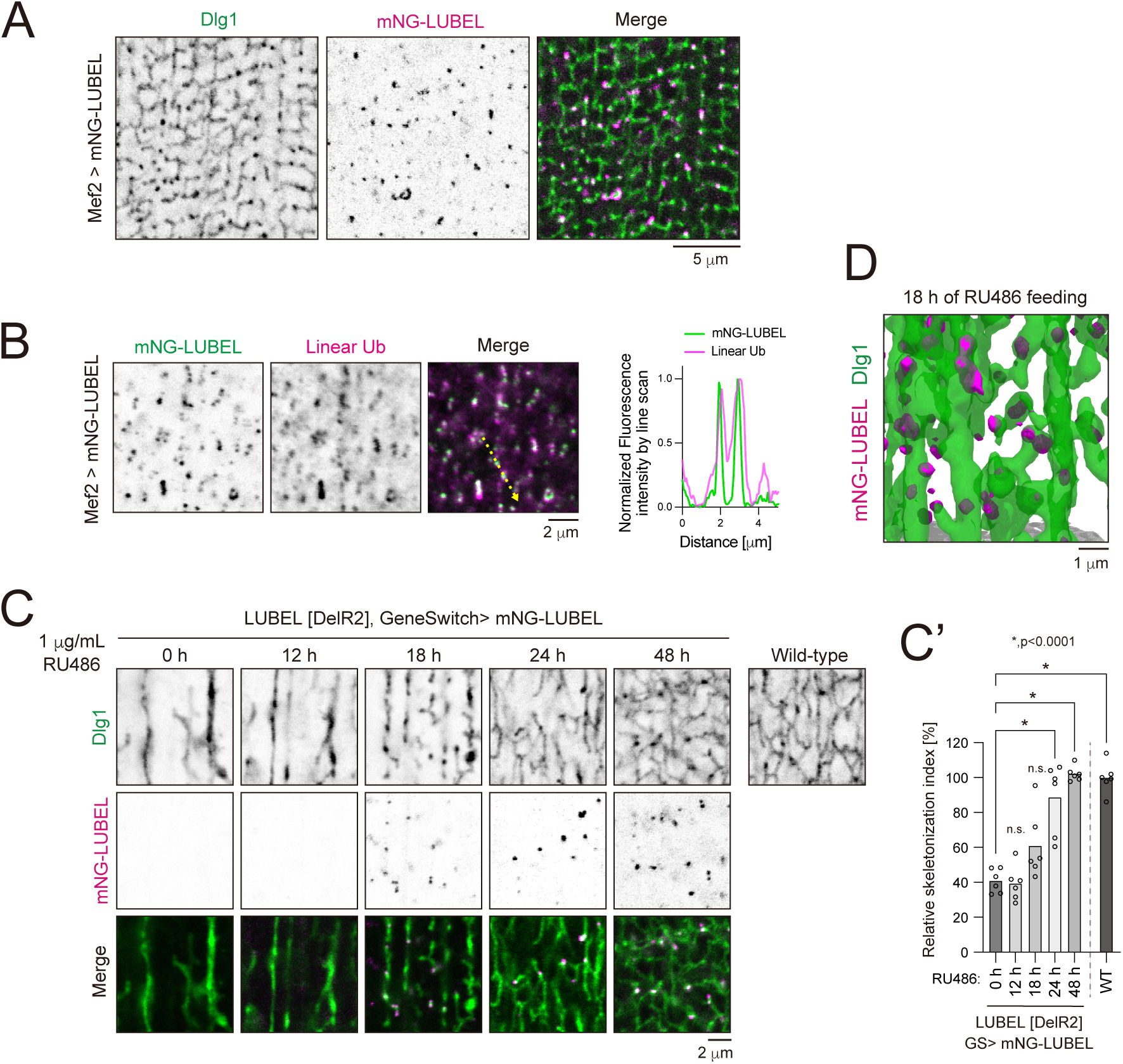
LUBEL and M1-linked ubiquitin chains form puncta on T-tubules. (A) Localization of mNG-LUBEL on T-tubules. 3IL BWMs expressing mNG-LUBEL were stained with anti-Dlg1 antibody. (B) Colocalization of mNG-LUBEL and M1-linked ubiquitin chains. 3IL BWMs expressing mNG-LUBEL were stained with anti-Dlg1 and anti-linear ubiquitin antibodies. (C, C’) LUBEL rescue experiment with temporally controlled mNG-LUBEL expression. mNG-LUBEL expression was regulated using the GeneSwitch system. Larvae were reared on RU486-containing food for the indicated duration before reaching the late 3IL stage. 3IL BWMs were stained with anti-Dlg1 antibody (C), and their relative skeletonization index was quantified (C’). (D) 3D reconstitution of z-series fluorescence images of mNG-LUBEL and Dlg1 in 3IL BWMs after 18 h of RU486 feeding.

After 48 hours of RU486 feeding, T-tubule defects in LUBEL mutants were almost completely rescued (Fig. 4C and C’), indicating that Dlg1-positive sheets transitioned into tubular structures upon mNG-LUBEL expression. Partial recovery was observed after 18 and 24 hours of RU486 feeding, suggesting that Dlg1-positive structures are forming T-tubules. Notably, mNG-LUBEL puncta were observed at the edge of these Dlg1-positive T-tubule intermediates. A 3D reconstruction of Z-series images (Fig. 4D) revealed that LUBEL puncta were frequently located at protrusions of forming T-tubules, suggesting a role in the early stage of T-tubule biogenesis.

### Condensation of LUBEL and M1-linked ubiquitin chains is essential for T-tubule formation

The spherical morphology of LUBEL and M1-linked ubiquitin chain-positive puncta suggests that they form protein condensates on membranes. Protein condensation is generally driven by multivalent interactions (*47*). Since LUBEL contains long IDRs, which can facilitate weak intra- and intermolecular interactions, we investigated whether LUBEL undergoes homotypic interactions. Myc-LUBEL^1-872^ was co-immunoprecipitated with HA-LUBEL^1-872^ (Fig. 5A). Furthermore, IP-MS of LUBEL^1-872^ identified LUBEL fragments corresponding to its C-terminal half (Fig. 3D and Data S3). These findings suggest that LUBEL interacts with itself intermolecularly through its IDRs (Fig. 5C).

**Figure 5.**
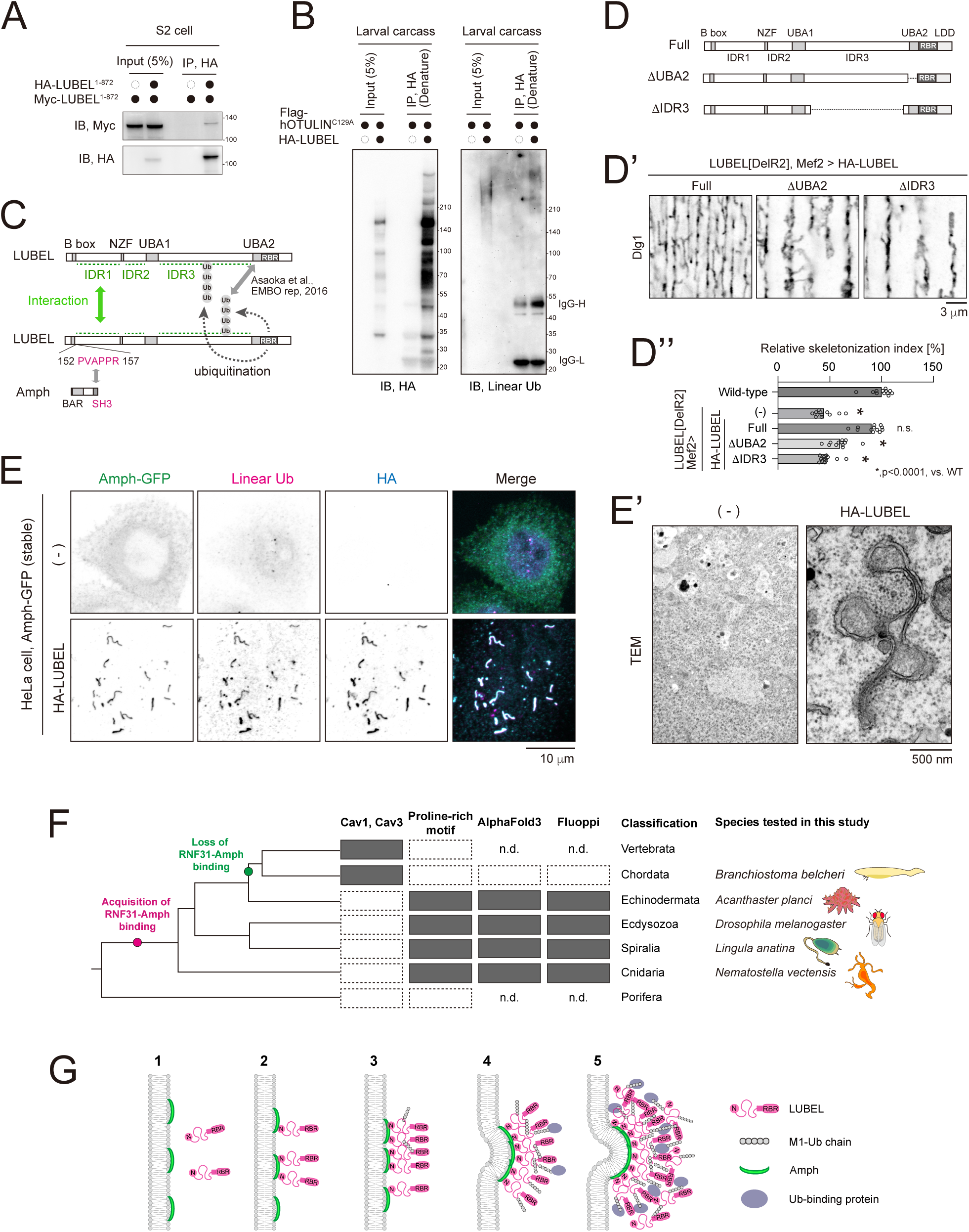
LUBEL condensate and membrane deformation. (A) Co-IP assay of the LUBEL N-terminal fragment. HA-LUBEL^1-872^ and Myc-LUBEL^1-872^ were co-expressed in S2 cells. Lysates were subjected to anti-HA IP and immunoblotted for the indicated antibodies. (B) LUBEL auto-ubiquitination. HA-LUBEL and the catalytically dead human OTULIN^C129A^ mutant were co-expressed in 3IL BWM. Carcass fillets were lysed in SDS-containing denaturing buffer, diluted 10-fold, and subjected to anti-HA IP. Samples were then immunoblotted with the indicated antibodies. (C) Schematic representation of LUBEL-mediated interactions. LUBEL potentially self-associates through IDRs and undergoes intra- or inter-molecular auto-ubiquitination. The UBA2 domain has an affinity for M1-linked ubiquitin chains. (D, D’, D’’) LUBEL rescue experiment using full-length (Full), ΔUBA2, or ΔIDR3 constructs (D). 3IL BWMs were stained with anti-Dlg1 antibody (D’), and the relative skeletonization index was quantified (D’’). (E, E’) LUBEL-dependent membrane tubulation in HeLa cells. HeLa cells stably expressing Amph-GFP were transfected with either HA-LUBEL or an empty vector and stained with the indicated antibodies (E) or analyzed by TEM (E’). (F) Evolutionary analysis of LUBEL-Amph interactions across animal species. For further details, see Fig. S4 and S5. (G) Schematic model of LUBEL-dependent T-tubule biogenesis.

Mammalian HOIP, a catalytic subunit of LUBAC, ubiquitinates itself and other subunits within the complex (*48*, *49*). By analogy, we hypothesized that LUBEL undergoes intra- and/or inter-molecular self-ubiquitination. To test this hypothesis, we performed denaturing IP using anti-HA antibody while co-expressing catalytically-dead human OTULIN to prevent the degradation of M1-linked ubiquitin chains (*30*, *49*). Immunoblotting with anti-M1-linked ubiquitin antibody revealed high-molecular-weight smear bands (Fig. 5B), suggesting that LUBEL undergoes self-ubiquitination. Since the UBA2 domain in LUBEL has been shown to interact with M1-linked ubiquitin chains (*30*), self-ubiquitination may contribute to the multivalency required for condensation (Fig. 5C).

To assess the functional significance of these interactions in T-tubule formation, we conducted rescue experiments using LUBEL constructs lacking either the UBA2 domain or the longest IDR (IDR3) (Fig. 5C and D). In stark contrast to the full-length protein, neither the UBA2-deleted nor IDR3-deleted constructs rescued T-tubule defects in LUBEL mutants (Fig. 5D’ and D’’). These results suggest that multivalent interactions mediated by the UBA2 and IDR3 in LUBEL are required for T-tubule biogenesis.

### LUBEL facilitates Amph-mediated membrane deformation

To evaluate whether LUBEL-mediated M1-linked ubiquitination is sufficient for Amph-dependent membrane deformation, we utilized HeLa cells as a semi-reconstituted system by constitutively expressing GFP-tagged *Drosophila* Amph (dmAmph). Stable expression of GFP-dmAmph alone did not induce membrane tubulation (Fig. 5E upper panels). Strikingly, the transient expression of LUBEL significantly enhanced the deformation of Amph-positive structures (Fig. 5E lower panels). Of note, LUBEL and M1-linked ubiquitin chains colocalized with the Amph-positive tubular structures. Furthermore, TEM analysis revealed irregular membrane structures in HeLa cells expressing LUBEL, which were absent in control cells (Fig. 5E’). These findings suggest that LUBEL-containing condensates are responsible for driving Amph-mediated membrane deformation.

### Evolutionary analysis of LUBEL-Amph interaction across animal species

The above results suggest that LUBEL condensate-mediated membrane deformation functions at the early stage of T-tubule biogenesis, similar to caveolins (*23*, *24*). Unlike in *Drosophila*, where T-tubule formation is independent of caveolins, Amph2-mediated T-tubule formation in mammals requires Cav3, which promotes the formation of membrane invaginations (*24*).

Caveolin is conserved across metazoans, including nematodes; however, not all caveolin family molecules contribute to membrane deformation (*26*, *50*). Caveolae-forming caveolins, Cav1 and Cav3, are conserved in chordates and vertebrates but absent in other metazoans (Fig. 5F, top).

Notably, *Drosophila* lacks caveolin genes entirely (*26*). Evolutionarily, T-tubules emerged before the caveolin-mediated membrane deformation mechanisms (*27–29*), indicating the presence of an alternative mechanism in invertebrates, including arthropods. For this reason, we speculate that LUBEL may be this alternative mechanism.

To explore this hypothesis, we conducted an evolutionary analysis of Amph and RNF31, the ortholog of LUBEL and HOIP, across animal species. Both Amph and RNF31 are widely conserved across metazoans. The proline-rich motif in RNF31 is present in a broad range of species, including those within Echinodermata, Ecdysozoa, Spiralia, and Cnidaria, but is absent in Vertebrata, Chordata, and Porifera (Fig. 5F). AlphaFold3 multimer modeling predicted that RNF31 interacts with Amph in *Acanthaster planci* (Ap), *Lingula anatine* (La), and *Nematostella vectensis* (Nv) but not in *Branchiostoma belcheri* (Bb) (Fig. S4). To experimentally validate this prediction, we employed the Fluoppi system, a fluorescence-based technology that detects protein-protein interaction via puncta formation in mammalian cultured cells. As anticipated, foci were observed in the combination of ApAmph-ApRNF31, LaAmph-LaRNF31, and NvAmph-NvRNF31, whereas foci were hardly detected for BbAmph-BbRNF31 (Fig. S5). These findings suggest that the RNF31-Amph interaction arose after the divergence from Porifera and was subsequently lost between the divergence of Echinodermata and the emergence of Vertebrata and Chordata (Fig. 5F). Notably, the evolutionary analysis indicates that the loss of the Amph-RNF31 interaction coincided with the acquisition of Cav1/3 (Fig. 5F). In stark contrast to the results in *Drosophila* (Fig. 1), neither RNF31 knockdown nor OTULIN overexpression significantly affected T-tubule morphology in zebrafish, suggesting that M1-linked ubiquitination by LUBAC is dispensable for T-tubule formation in vertebrates. Based on these findings, we propose that LUBEL-mediated M1-linked ubiquitination plays a role analogous to caveolin in T-tubule biogenesis despite significant differences in their underlying molecular mechanisms.

## Discussion

In this study, we discovered an unexpected role for M1-linked ubiquitination in T-tubule biogenesis. Based on our findings, we propose the following mechanism (Fig. 5G): (1) Amph initially localizes to the membrane via its amphipathic helix. (2) LUBEL is recruited to the membrane through the interaction between the SH3 domain of Amph and the proline-rich motif (152–157) in LUBEL. (3) LUBEL ubiquitinates itself and its proximity molecules. (4) Multivalent interactions mediated by the IDR and UBA domain in LUBEL promote the formation of protein condensates on the Amph-positive membrane. (5) These condensates facilitate membrane tubulation by increasing the local concentration of Amph and/or inducing subtle deformations in membranes.

Membrane localization of Amph is not sufficient to drive membrane tubule extension (Fig. 2). Previous studies have shown that while the amphipathic helix of Amph mediates initial membrane binding, tubulation requires high protein densities that enable the formation of oligomeric BAR domain scaffolds (*51*). Given that a minimum threshold of Amph density is necessary for extensive tubulation and that LUBEL condensates are essential for Amph-mediated tubulation by directly interacting with Amph (Fig. 3), it is likely that LUBEL-containing condensates cluster multiple Amph molecules, increase their local concentration on the membrane, and thereby induce membrane deformation.

The initial stage of membrane deformation, where a flat membrane transforms into a tubular shape, is expected to have the highest energy barrier. LUBEL condensates may induce subtle deformations in membranes that stimulate Amph polymerization, in addition to increasing Amph concentration, acting as a structural template. Furthermore, molecular crowding and viscoelastic effects are plausible mechanisms through which protein condensates generate positive membrane curvature. Our preliminary FRAP data on mNG-LUBEL suggest that LUBEL-containing condensates exhibit low fluidity. Therefore, we propose that molecular crowding plays a key role in the membrane deformation mediated by LUBEL condensates (Fig. 5F). Of note, Amph concentration and subtle membrane deformation by condensates are not mutually exclusive— both mechanisms may synergistically contribute to the early stages of T-tubule biogenesis, facilitating further Amph polymerization to drive membrane tubule extension.

Our findings suggest that the LUBEL/RNF31 condensate-mediated mechanism represents a caveolin-independent alternative for Amph-mediated tubulation initiation (Fig. S6). Evolutionary analysis suggests that the RNF31-dependent membrane deformation mechanism emerged early in animal evolution but was subsequently lost in specific lineages (Fig. 5F). Notably, the loss of the Amph-RNF31 interaction coincided with the emergence of Cav1/3 (Fig. 5F), suggesting a functional transition. Based on these findings, we propose that the ancestral RNF31-dependent mechanism was replaced by the caveolin-dependent one during evolution. In vertebrates, the functional specialization of M1-linked ubiquitination in immune regulation may have taken precedence, possibly reflecting the increasing complexity of their immune systems. In contrast, invertebrates have largely retained the RNF31-Amph interaction primarily for its role in membrane deformation processes, such as T-tubule formation.

A ubiquitin reader that recognizes M1-linked ubiquitin chains may play a role in LUBEL-mediated T-tubule biogenesis, similar to other ubiquitin-dependent processes (*52*). However, apart from LUBEL, no ubiquitin-binding domain-containing factors were identified in our RNAi screening (Fig. 1D and Data S2). The ubiquitin-binding UBAN domain, found in mammalian ABIN-1/2/3 and NEMO, is known for its high affinity for M1-linked ubiquitin chains and is also present in Optineurin (*53*). Additionally, M1-linked ubiquitin-binding NZF and zinc finger domains have been identified in HOIL-1 and A20, respectively (*53*). In contrast, Kenny is currently the only recognized M1-linked ubiquitin-binding protein in *Drosophila* (*42*); however, our results indicate that it is dispensable for T-tubule biogenesis (Fig. S2). Further investigation is needed to determine if a reader associates with the M1-linked ubiquitin chains involved in T-tubule biogenesis.

Structurally, the M1-linked ubiquitin chain resembles the K63-linked chain (*54*). It would be interesting to test whether an artificial LUBEL construct, in which the RBR is replaced by the RING domain of TRAF2 or TRAF6, could function in T-tubule biogenesis. Testing this construct would clarify whether the M1-linked ubiquitin chain can substitute for the K63-linked chain in this process. Furthermore, we cannot rule out the possibility that additional substrates beyond LUBEL are M1-linked ubiquitinated to facilitate T-tubule biogenesis. Comprehensive identification of such substrates in *Drosophila* will be crucial. Recent advances in ubiquitin-specific proximity-labeling technology offer promising solutions to achieve this goal (*55*, *56*).

Our data demonstrates the novel concept that ubiquitin-mediated condensates regulate BAR domain proteins. A significant subset of BAR domain proteins contain scaffolding domains, including SH2, SH3, PDZ, and WW domains (*1*), which can interact with ubiquitin E3 ligases (*20*, *57*, *58*). We propose that, beyond M1-linked ubiquitin chains, other ubiquitin linkage types may also regulate BAR protein functions, broadening our understanding of the molecular mechanisms underlying membrane deformation.

## Materials and methods

### Reagents and antibodies

Streptavidin-HRP (#S911), Phalloidin-Alexa546 (#A22283), anti-HA magnetic beads (#88837), and glutathione agarose (#16100) were purchased from Thermo Fisher Scientific (Waltham, MA, USA). Anti-Dlg1 antibody (clone 4F3, RRID: AB_528203) was obtained from the Developmental Studies Hybridoma Bank (Iowa City, IA, USA). Anti-*Drosophila* Amph antibody was generously provided by Cahir O’Kane (*17*). Anti-myc (#562), anti-HA (#561), and anti-V5 (#PM003) antibodies were purchased from MBL (Nagoya, Japan). Anti-HA antibody (clone 3F10, #11867423001) was obtained from Roche (Basel, Switzerland). Anti-M1-linked ubiquitin antibody was kindly provided by Genentech (South San Francisco, CA, USA) (*59*).

### Generation of transgenic flies

cDNAs encoding the LUBEL and OTULIN variants were integrated into the pUAST-attB vector using standard molecular cloning techniques. LUBEL and human OTULIN cDNAs were provided by Fumiyo Ikeda (*30*). The Amph isoform C cDNA was amplified from larval carcass fillets. Microinjections into the attP landing site stocks were performed by Wellgenetics Inc. (Taipei, Taiwan).

### *Drosophila* strains and maintenance

Flies were maintained under standard conditions at 25°C unless otherwise stated. The *w^1118^*strain was used as a control for LUBEL mutant strains. For muscle-targeted gene expression, Mef2-GAL4 was used. UAS-*LacZ* was used as a control for UAS transgenes. Detailed genotypes are described in Table S1. The following fly stocks were used:

(1) *y,w*; *P*{*w^+mC^=GAL4-Mef2.R*}3 (Bloomington *Drosophila* Stock Center, BDSC_27390), referred to as DMef2-GAL4);
(2)*w*; UAS-V5-miniTurbo (kindly provided by Steve Jean, Univ. of Sherbrooke), referred to as UAS-V5-miniTurbo;
(3)*w*; *{UAS-Amph^isoform_A^-miniTurbo}attP40* (this study), referred to as UAS-Amph-V5-miniTurbo;
(4)*w; P{w[+mC]=UAS-Dcr-2.D}10* (BDSC_24651), referred to as UAS-Dcr2;
(5)*w*, *P{PTT-GC}dlg1[YC0005]* (BDSC_50859), referred to as Dlg1-GFP;
(6)*w*; *P*{*w^+mC^=UAS-lacZ.B*}*Bg4-1-2* (BDSC_1776), referred to as UAS-LacZ;
(7)*y,v; P{TRiP.JF02883}attP2* (BDSC_28048), referred to as UAS-IR-Amph;
(8)*w*, *UAS-IR-LUBEL^11321R-2^* (NIG-RNAi_11321R-2), referred to as UAS-IR-LUBEL_NIG;
(9)*w*; *UAS-IR-LUBEL^KK106140^* (VDRC_100651), referred to as UAS-IR-LUBEL_KK;
(10)*w*; *UAS-IR-LUBEL^GD18055^* (VDRC_18055), referred to as UAS-IR-LUBEL_GD;
(11)*y,w; Mi{ET1}LUBEL[MB00197]* (BL_22725), referred to as LUBEL^MI^;
(12)*LUBEL^DelR2^* (30);
(13)*LUBEL^CC/SS^* (30);
(14)*w*; *{UAS-3xFLAG-hsOTULIN}attP2* (this study), referred to as UAS-hOTULIN;
(15)*w*; *{UAS-3xFLAG-hsOTULIN^C129A^}attP2* (this study), referred to as UAS-hOTULIN^C129A^;
(16)*w; Rel[E20] e[s]* (BDSC_9457), referred to as Rel[E20];
(17)*w; PBac{w[+mC]=PB}key[c02831]* (BDSC_11044), referred to as kenny[c02831];
(18)*w*; *{UAS-3xHA-LUBEL}attP2* (this study), referred to as UAS-HA-LUBEL;
(19)*w*; *{UAS-3xHA-LUBEL^1-872^}attP2* (this study), referred to as UAS-HA-LUBEL^1-872^;
(20)*w*; *{UAS-3xHA-LUBEL^872-2902^}attP2* (this study), referred to as UAS-HA-LUBEL^872-2902^;
(21)*w*; *{UAS-miniTurbo-3xHA-LUBEL}attP2* (this study), referred to as UAS-mnTb-LUBEL;
(22)*w*; *{UAS-mNeonGreen-3xHA-LUBEL}attP2* (this study), referred to as UAS-mNG-LUBEL;
(23)*w*; *{UAS-3xHA-LUBEL^112466-2521^}attP2* (this study), referred to as UAS-HA-LUBEL^11UBA2^;
(24)*w*; *{UAS-3xHA-LUBEL ^Δ1214-2441^}attP2* (this study), referred to as UAS-HA-LUBEL^11IDR^;
(25)*w*; *{UAS-Amph^isoform_C^:3xV5}attP2* (this study), referred to as UAS-Amph-V5;

### RNAi screening

UAS-RNAi lines (Data S2) were crossed with *w, Dlg1-GFP*; *Mef2-GAL4, UAS-Dcr2*. The detailed procedures were previously reported (*32*). Briefly, immobilized 3ILs were mounted between a slide glass and a cover glass following the protocol described by Zitserman and Roegiers (*60*). Dlg1-GFP signals in BWM were imaged using the confocal microscope FV3000 (EVIDENT, Tokyo, Japan) through the dorsal cuticle.

### Immunostaining of LBWMs

Third-instar larvae were dissected in dissection buffer (5 mM HEPES, 128 mM NaCl, 2 mM KCl, 4 mM MgCl₂, 36 mM sucrose) on a 35 mm dish coated with silicone elastomer (SYLGARD™ 184, Dow Corning, #3097358-1004, Midland, MI, USA) using scissors, forceps, and micro pins (Fine Science Tools, #11295-10, #15000-08, #26002-10, Foster City, CA, USA). The larval carcass fillets were fixed with 4% paraformaldehyde (PFA) in phosphate buffer (Nacalai Tesque, #09154-85, Kyoto, Japan) for 20 min at room temperature. The fixed fillets were washed with PBS three times and incubated with the primary antibody diluted in blocking buffer (0.3% BSA, 0.6% Triton X-100 in PBS) overnight at room temperature. The fillets were washed with PBS three times and incubated with the secondary antibody diluted in a blocking buffer for 1 hour at room temperature. The fillets were then washed with PBS three times and mounted with FluorSave (Millipore/Merck, Burlington, Massachusetts, USA).

### Confocal fluorescence microscopy

Both live and fixed samples were observed using a confocal microscope FV3000 (EVIDENT, Tokyo, Japan) equipped with a 60× silicone/1.30 NA objective lens (EVIDENT, Tokyo, Japan). FLUOVIEW (EVIDENT, Tokyo, Japan) was used for image acquisition, and the exported images were processed and analyzed with ImageJ (NIH, Bethesda, MD, USA).

### Proximity labeling in LBWMs and mass spectrometry analysis of biotinylated peptides

Second or early third instar larvae expressing miniTurbo-fused constructs under the control of Mef2-GAL4 were reared on fly media supplemented with 1 mM biotin for 2 days at 25°C. The biotin-fed 3IL were then dissected to obtain carcass fillets containing BWMs. The fillets were lysed using a guanidine buffer composed of 6 M guanidine hydrochloride, 20 mM Tris-HCl, 10 mM Tris(2-carboxyethyl)phosphine hydrochloride, 40 mM chloroacetamide, and a protease inhibitor cocktail (Nacalai Tesque, #04080-11). The biotinylated peptides were purified as described previously (*34*). Briefly, the extracted protein solution was digested with trypsin (MS grade, Thermo Fisher Scientific), and biotinylated peptides were enriched using MagCapture HP Tamavidin 2-REV magnetic beads (#133-18611, FUJIFILM Wako, Japan). The resulting samples were then proceeded for mass spectrometry.

### Mammalian cell culture

HeLa and Plat-E cells were cultured in Dulbecco’s Modified Eagle Medium (DMEM, Nacalai Tesuque, #08458-45) supplemented with 10% fetal bovine serum (FBS) and 1% penicillin-streptomycin. Cells were maintained at 37°C in a humidified environment with 5% CO₂.

### *Drosophila* S2 cell culture

S2 cells were cultured at 25°C in Schneider’s *Drosophila* medium (Thermo Fisher Scientific, #21720-024) supplemented with 10% FBS and penicillin-streptomycin.

### Plasmid transfection

JetPRIME Transfection Reagent (Polyplus-transfection, Illkirch, France) was used for plasmid transfection into HeLa, Plat-E, and S2 cells. Cells were plated in wells or dishes one day before transfection to achieve 50-70% confluence on the day of transfection. Plasmid DNA was diluted using JetPRIME buffer (1 μg DNA per 100 μL buffer). Following this, JetPRIME Transfection Reagent was added to the diluted DNA at a ratio of 2:1 (2 μL of JetPRIME for 1 μg of plasmid DNA) and incubated at room temperature for 10 min. The resulting DNA/JetPRIME complexes were then added dropwise to cells in growth medium, and the transfected cells were incubated for 2 days to allow for gene expression.

### Generation of HeLa cells stably expressing dmAmph-EGFP

Plat-E cells were transfected with pMRX-Amph_C-GFP-IRES2-puroR and pVSV-G. The supernatants containing recombinant retrovirus were filtered through a 20 µm filter. HeLa cells were then infected with the retrovirus solution, supplemented with 8 μg/mL polybrene for 6 hours. Stable transformants were then selected using 1 μg/mL puromycin for one week.

### Immunostaining of HeLa cells

HeLa cells were cultured on 15 mm cover glasses in 12-well plates and fixed with 4% PFA in PBS for 10 min at room temperature. The fixed cells were washed with PBS twice and incubated with the primary antibody diluted in blocking buffer (1% BSA, 0.1% Triton X-100 in PBS) for 1 hour at room temperature. The cells were washed twice again with PBS and incubated with the secondary antibody diluted in a blocking buffer for 1 hour at room temperature. The stained samples were then washed twice with PBS and mounted using FluorSave (Millipore/Merck, Burlington, MA, USA).

### Protein-protein interaction assay using Fluoppi system

The coding region of the Amph SH3 domains and the RNF31 proline-rich regions from *Branchiostoma belcheri*, *Acanthaster planci*, *Lingula anatina*, and *Nematostella vectensis* were synthesized by Integrated DNA Technologies, Inc. (Coralville, IA, USA). These synthesized DNA fragments were cloned into the pAzamiGreen and pHA-Ash vectors using the *Eco*RI and *Bam*HI restriction sites. HeLa cells were transfected with pAzamiGreen-RNF31 (proline-rich region) and pHA-Ash-Amph (SH3 domain), followed by immunostaining with anti-HA antibody (3F10). Confocal imaging was performed using a FV3000 microscope (EVIDENT, Tokyo, Japan), and image analysis was conducted using ImageJ Fiji. Each cell was manually cropped and processed using a median filter and the “Subtract Background” function. The resulting single-cell images were converted to binary images using the “Threshold” function. To assess colocalization, binary images from AzamiGreen and HA-Ash channels were multiplied using the ImageCalculator function. The percentage of colocalized dots was then quantified using the “Measure” function. For the Fluoppi analysis (Fig. S5), a minimum of 15 cells were analyzed for each combination.

### Immunoprecipitation and GST pulldown

S2 cells grown in 6-well plates or 20 third-instar larval carcass fillets were lysed with 200 μL of lysis buffer (1% NP-40, 1 mM EDTA, 20 mM Tris-HCl pH 7.4, 150 mM NaCl) and centrifuged at 21,130 g for 10 min at 4°C. For input samples, 20 μL of the supernatant was mixed with 20 μL of 2x sample buffer (4% SDS, 20% glycerol, 125 mM Tris-HCl pH 6.8, 0.001% bromophenol blue, 200 mM dithiothreitol). For immunoprecipitation, 150 μL of the supernatant was incubated with 10 μL of anti-HA magnetic beads (Thermo Fisher Scientific, #88837) or GST-fused protein-conjugated glutathione agarose (Thermo Fisher Scientific, #16100) for 90 min at 4°C. The beads were then washed three times with 500 μL of lysis buffer and eluted with 20 μL of 1x sample buffer by heating at 95°C for 5 min. For mass spectrometric analysis, 5 μL of anti-HA magnetic beads were used for immunoprecipitation. The samples were then washed and resuspended in 50 mM triethylammonium hydrogen carbonate solution (Fujifilm Wako Chemicals, Osaka, Japan, #206-08381)

### Western blotting

The samples were subjected to SDS-PAGE using a precast gel (SuperSep Ace, 5-20%, 17 wells, Fujifilm Wako Chemicals, #194-1521). Proteins in the acrylamide gel were transferred onto a PVDF membrane (Millipore/Merck, Immobilon-PSQ, #ISEQ00010) using transfer buffer (0.1% SDS, 1.2% Tris base, 1.44% glycine, 20% methanol). The transfer was conducted at a current of 2 A or a voltage of 15 V for 1 hour with the Trans-Blot Turbo transfer system (BioRad, #1704150J1). Following the transfer, the membrane was blocked (1% BSA, 0.1% Tween-20, 10 mM Tris-HCl, pH 8.0) for 10 min at room temperature. It was then incubated with the primary antibody diluted in Can Get Signal buffer (Toyobo, #NKB-101, Osaka, Japan) overnight at room temperature. The membrane was washed three times with TBS-T (0.1% Tween-20, 10 mM Tris-HCl, pH 8.0, 150 mM NaCl) and then incubated with an HRP- or StarBright Blue 700-conjugated secondary antibody diluted in blocking buffer for 1 h at room temperature. The membrane was washed again three times with TBS-T. Immunoreactive bands were detected using Clarity Western ECL Substrate (Bio-Rad, Hercules, CA, USA) and ChemiDoc MP Imaging System (Bio-Rad, Hercules, CA, USA). The resulting images were processed using ImageJ.

### Mass spectrometry analysis of co-immunoprecipitated proteins

HA-LUBEL^1-872^ or LacZ (control) was expressed under the control of Mef2-GAL4, and the progenies were raised at 25°C. The 3IL carcass fillet lysates were subjected to anti-HA immunoprecipitation. The immunoprecipitated proteins bound to the beads were reduced and alkylated with 10 mM TCEP, 40 mM CAA, and 5% SDS at 65°C for 10 min. Proteins in the supernatants were collected and cleaned up using the SP3 method (*61*) and the ‘on-beads’ digestion to peptides was performed using trypsin for 18 hours at 37°C. Peptides were further purified by SP3 and eluted from the beads by 2% DMSO. Peptides were analyzed by an EASY-nLC 1200 UHPLC-coupled Orbitrap Fusion mass spectrometer (Thermo Fisher Scientific). The peptides in 0.1% TFA were separated on a C18 reversed-phase column (75 μm × 150 mm; Nikkyo Technos) with a linear 4-32% acetonitrile (ACN) gradient for 0-100 min with a flow rate of 300 nL/min, followed by an increase to 80% ACN for 10 min and a final hold at 80% ACN for 10 min. Peptides were ionized by electrospray ionization at 2.0 kV. The data-dependent acquisition method was used to acquire MS/MS spectra with a maximum duty cycle of 3 s. The primary mass spectrometry scan (MS1) was acquired in the Orbitrap at a 120,000 resolution with a maximum injection time of 50 ms. Then, the most abundant ions within a scan range of 375-1,500 m/z based on the m/z signal with charge states of 2-7 from that scan were chosen from the MS1 for collision-induced dissociation in the HCD cell and MS2 analysis in the linear ion trap with an AGC target of 1e4, an isolation window of 1.6 m/z, a maximum injection time of 35 ms, and a normalized collision energy of 30. Dynamic exclusion was set to 20 s. Raw data were directly analyzed against the SwissProt database restricted to *Drosophila melanogaster* using Proteome Discoverer 2.5 (Thermo Fisher Scientific) with the Sequest HT search engine. The search parameters were as follows: trypsin as an enzyme with up to two missed cleavage, precursor mass tolerance of 10 ppm, fragment mass tolerance of 0.6 Da, carbamidomethylation of cysteine as a fixed modification, and acetylation of the protein N-terminus and oxidation of methionine as variable modifications. Peptides were filtered at a false discovery rate of 1% using the Percolator node. Label-free quantification was performed based on the intensities of precursor ions using the Precursor Ions Quantifier node.

### TEM and FIB-SEM

Third instar larvae (3IL) were pinned on a silicone elastomer-covered Petri dish and dissected directly in a fixative solution containing 2% PFA, 2.5% glutaraldehyde, 150 mM sodium cacodylate, and 5 mM calcium chloride (pH 7.4). The larval carcass fillets were fixed for 2 hours at room temperature and then overnight at 4°C. The dissected fillets were washed with 0.1 M phosphate buffer (pH 7.4), post-fixed in 1% osmium tetroxide (OsO₄) buffered with 0.1 M phosphate buffer for 2 hours, dehydrated through a graded ethanol series, and embedded flat in Epon 812 (TAAB, Aldermaston, UK).

For TEM of BWMs, ultrathin sections (70 nm thick) were mounted on copper grids coated with Formvar (Nisshin EM, Tokyo, Japan) and double-stained with uranyl acetate and lead citrate. The samples were then observed using a JEM-1400Flash transmission electron microscope (JEOL, Tokyo, Japan).

FIB-SEM tomography was performed using a Helios NanoLab 660 FIB-SEM (Thermo Fisher Scientific, Waltham, MA, USA) as previously described (*62*). Briefly, serial FIB-SEM images were acquired using Auto Slice and View imaging software (Thermo Fisher Scientific) on the same instrument. Two sets of 500 serial images at 50 nm depth intervals were obtained from wild-type and LUBEL mutant BWMs using a backscattered electron detector at an acceleration voltage of 3.0 kV. After alignment, individual FIB-SEM images were manually segmented to identify target membrane structures using AMIRA 6.1 reconstruction software (Thermo Fisher Scientific), followed by 3D reconstruction.

For TEM of HeLa cells, a glass coverslip with a grid (GC1300, Matsunami Glass, Osaka, Japan) was coated with carbon using a vacuum evaporator IB-29510VET (JEOL, Tokyo, Japan). HeLa cells stably expressing dmAmph-GFP were cultured on the coated coverslip. The cells were then transfected with either an empty vector or plasmid encoding HA-LUBEL and incubated for 16 hours. Subsequently, the cells were fixed with 2% PFA and 0.5% glutaraldehyde in phosphate buffer (pH 7.4) for 1 hour at room temperature. The positions of HA-LUBEL-transfected cells were identified by confocal microscope FV3000 (EVIDENT, Tokyo, Japan). After fluorescence microscopy, the cells were further fixed with 2.5% glutaraldehyde in 0.1 M phosphate buffer (pH7.4) for 2 hours at 4°C, followed by post-fixation in 1% osmium tetroxide (OsO₄) in 0.1 M phosphate buffer for 2 hours. The samples were then dehydrated through a graded ethanol series and embedded flat in Epon 812 (TAAB, Aldermaston, UK). Ultrathin sections were prepared by trimming the same region observed by confocal microscopy. These sections, with a thickness of 70 nm, were mounted on copper grids, double-stained with uranyl acetate and lead citrate, and observed using a JEM-1400Flash transmission electron microscope (JEOL, Tokyo, Japan).

### Prediction of RNF31-Amph interactions using AlphaFold

Protein structure prediction for *Drosophila melanogaster* (Dm) in Fig. 3G was performed using AlphaFold2 via ColabFold (v1.5.2) on Google Colaboratory (*63*). The publicly available notebook, “AlphaFold2.ipynb”, was used. The multiple sequence alignments were generated using MMseqs2 against the UniRef90 and MGnify databases. Structure prediction was performed using the AlphaFold2_multimer_v3 model with five ensemble predictions.

For the analysis shown in Fig. S4, including Dm, structure prediction was performed using AlphaFold3 (*64*) via the AlphaFold server (https://golgi.sandbox.google.com). The protein sequences used for structure prediction were obtained from publicly available databases. The protein sequences of *Acanthaster planci* (Ap), *Lingula anatine* (La), and *Nematostella vectensis* (Nv), and *Branchiostoma belcheri* (Bb) were identified via NCBI BLASTp using the protein sequence of Dm as the query. The identifiers of the protein sequences are as follows: DmLUBEL/RNF31 (ID: Q8IPJ3), DmAmph (ID: Q7KLE5), ApRNF31 (ID: XP_022084012.1), ApAmph (ID: XP_022084611.1), LaRNF31 (ID: XP_013410818.1), LaAmph (ID: XP_013391361.1), NvRNF31 (ID: XP_032219182.2), NvAmph (ID: XP_048576432.1), BbRNF31 (ID: XP_019627184.1), and BbAmph (ID: XP_019613513.1).

### Image analyses

For quantification of T-tubules, the middle sections of LBWM were imaged using confocal microscopy. Each image had a resolution of 1024 × 1024 pixels, corresponding to a physical size of 70.71 μm. Prior to analysis, regions surrounding the muscle cells were removed. Image processing and quantification were conducted using ImageJ Fiji (NIH, Bethesda, MD, USA). To minimize noise and background, a median filter background subtraction was applied to each image. The processed images were then converted to binary format and further processed using the “Skeletonize” function. The skeletonization index was defined as the mean intensity of the skeletonized binary images. The ImageJ macro used for this analysis is provided below:

~~~
showMessage(”Select the folder containing Anti-Dlg1 images”);
openDir = getDirectory(”Choose a Directory”);
list = getFileList(openDir);
Array.show(list);
for (i=0; i<list.length; i++){
 open(openDir+list[i]);
 run(“Median…”, “radius=2”);
 run(“Subtract Background…”, “rolling=20”);
 setAutoThreshold(”Huang dark”);
 run(”Convert to Mask”);
 run(“Skeletonize”);
 run(”Measure”); close();
}
~~~

For the skeletonization analysis (Fig. 1F, 1G, 3B, 3C, 3H, 4C, 5D, and S2), at least six images from three or more animals were analyzed for each genotype and time point.

For line plot analysis, fluorescence intensity profiles along the selected line were obtained using the “Plot Profile” function in ImageJ Fiji (NIH). The intensity data were normalized in Microsoft Excel by setting the maximum and minimum intensities to 1 and 0, respectively. Graphs were generated using GraphPad Prism (GraphPad Software, Boston, MA, USA).

### Statistical analysis

GraphPad Prism 9 software (GraphPad, Prism9 version 9.5.1) was used for statistical analyses. When two genotypes were used in an experiment, Student’s t-test was used. When more than two genotypes were used in an experiment, one-way ANOVA with Dunnett’s multiple comparisons test was used. P<0.05 was regarded as statistically significant.

## Acknowledgments

We are grateful to A. Kiger (UCSD), Cahir O’Kane (Univ. of Cambridge), BDSC, VDRC, NIG-FLY, and Genentech Inc. for reagents; T. Murakawa (Science Tokyo), T. Matsuyama (Science Tokyo), S. Kanamaru (Science Tokyo), W. Kojima (Science Tokyo), T. Kotani (Science Tokyo), and T. Miyaki (Juntendo Univ.) for their technical assistance; T. Fujimoto (Juntendo Univ.), N. Matsuda (Science Tokyo), Y. Yoshida (TMiMS), F. Tokunaga (OMU), D. Oikawa (OMU) T. Nishimura (Univ. of Osaka), and T. Noda (Univ. of Osaka) for helpful discussion; M. Landekic (McGill Univ.) for English editing; the Biomaterials Analysis Division of the Institute of Science Tokyo for the DNA sequencing.

## Funding

This work was supported in part by Grant-in-Aid for Transformative Research Areas (B) (grant number 21H05147, NF), Japan Science and Technology Agency (JST) PRESTO (grant number JPMJPR18H8, NF), and Grant-in-Aid for Scientific Research (B) from the MEXT (grant number 25K02270, NF).

## Author contributions

Conceptualization: NF

Methodology: KK, YH, HY, FI, HK, NF

Investigation: KK, YH, HY, KN, KM, TN, MK, YS. KA, SK, KI, HK, NF

Visualization: KK, YH, HY, KN, NF

Funding acquisition: NF

Project administration: NF Supervision: NF

Writing – original draft: KK, NF

Writing – review & editing: KK, YH, HY, SK, FI, HK, NF

## Competing interests

Authors declare that they have no competing interests.

## Supplementary Materials

Figs. S1 to S6

Tables S1 Data S1 to S4

Movies S1 and S2

## Supplementary material legends

**Figure S1. LUBEL RNAi phenocopies Amph RNAi in T-tubule morphology.** (A) Localization of Amph-mnTb or mnTb. Images of anti-Dlg1 and anti-V5 staining in 3IL BWMs. (B) LUBEL or Amph RNAi on Dlg1-GFP-positive T-tubule morphology in 3IL BWMs. Images were acquired through the cuticle using a confocal microscope.

**Figure S2. T-tubule formation is independent of the NF-κB signaling pathway.** Images showing anti-Dlg1 staining of 3IL BWMs in the indicated genotypes with the relative skeletonization index quantified.

**Figure S3. Characterization of the interaction between LUBEL and Amph.** (A) Localization of HA-LUBEL^1-872^. 3IL BWMs expressing HA-LUBEL^1-872^ were stained with anti-Dlg1 and anti-HA antibodies. (B) Schematic representation of LUBEL and Amph truncations. (C) GST pulldown assay of truncated Amph and LUBEL constructs. Myc-LUBEL^1-872^ was expressed in S2 cells. Lysates containing Myc-LUBEL^1-872^ were incubated with beads conjugated to GST-fused truncated Amph isoform C (GST-BAR and GST-SH3). The resultant samples were immunoblotted with anti-Myc antibody or stained with CBB. (D) Structure of the Amph SH3 and LUBEL-N fragment complex using AlphaFold 2. Four proline-rich motifs and a B-box zinc finger are shown. (E) Schematic representation of LUBEL point mutants. (E’) Co-IP assay of the LUBEL point mutants and Amph. The Myc-LUBEL^1-872^ fragments harboring the point mutations indicated in (E) were co-expressed with Amph isoform C-HA in S2 cells. Lysates were subjected to anti-Myc IP and immunoblotted for anti-Myc and anti-HA antibodies.

**Figure S4. AlphaFold prediction of the RNF31-Amph interaction.** Predicted structures of the Amph SH3–RNF31 complex for each species shown in Figure 5F, generated using AlphaFold3. The Amph SH3 domain is depicted with its surface electrostatic potential, while RNF31/LUBEL is represented as cyan stick models. Below each structure, the domain organization of RNF31/LUBEL from each species is shown, with the Amph-interacting region (proline-rich motif) highlighted in cyan, along with its position and sequence.

**Figure S5. Protein-protein interaction assay using Fluoppi system.** (A-E) Fluoppi analysis of the interaction between RNF31 and Amph in HeLa cells. HeLa cells transiently expressing the indicated constructs from each species listed were stained with anti-HA antibodies. (F) Quantification of foci positive for both RNF31 and Amph.

**Figure S6. Schematic models of the early stages of T-tubule biogenesis in mammals and *Drosophila.***

**Table S1. Detailed Drosophila genotypes used in each figure**

Detailed genotypes, stock center identifiers, references, and temperatures are provided.

**Data S1. Proximity proteomics of Amph-mnTb**

Output table generated by Proteome Discoverer software.

**Data S2. RNAi screening of T-tubule formation**

Stock center identifiers, CG numbers, and result summaries are shown.

**Data S3. IP-MS of HA-LUBEL^1-872^**

Output table generated by Proteome Discoverer software.

**Data S4. Proximity proteomics of mnTb-LUBEL**

Output table generated by Proteome Discoverer software.

**Movie S1**

Segmentation and 3D reconstitutions of T-tubules in wild-type larval BWMs, derived from FIB-SEM stacks.

**Movie S2**

Segmentation and 3D reconstitutions of T-tubule-related membrane structures in *LUBEL^CC/SS^* mutant larval BWMs, derived from FIB-SEM stacks.

## Notes

### Competing Interest Statement

The authors have declared no competing interest.

