## Supplementary figures for "Linear ubiquitination triggers Amph-mediated T-tubule biogenesis"

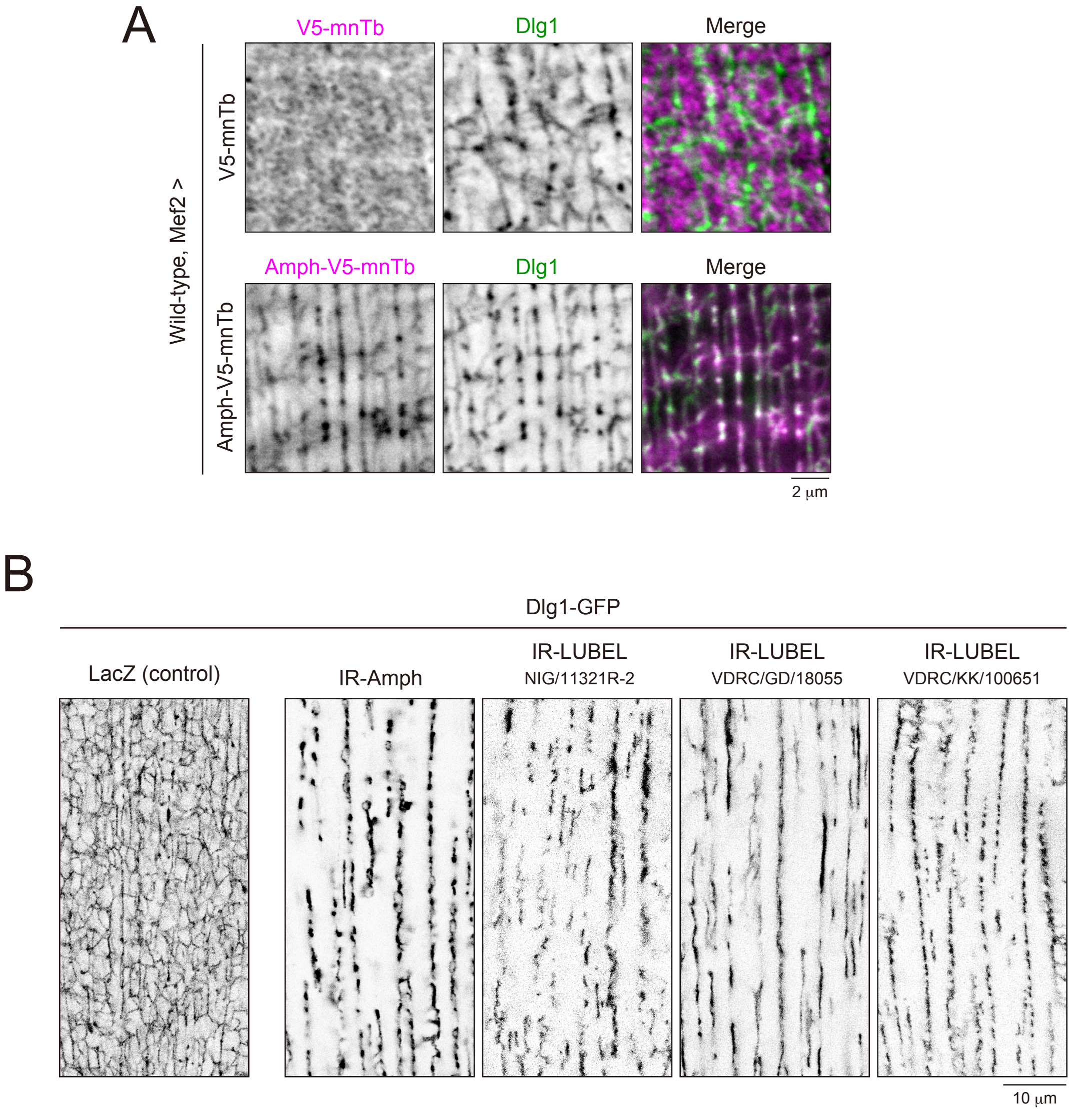


**Figure S1. LUBEL RNAi phenocopies Amph RNAi in T-tubule morphology**
(A) Localization of Amph-mnTb or mnTb. Images of anti-Dlg1 and anti-V5 staining in 3IL BWMs. (B) LUBEL or Amph RNAi on Dlg1-GFP-positive T-tubule morphology in 3IL BWMs. Images were acquired through the cuticle using a confocal microscope.

### Figure S1

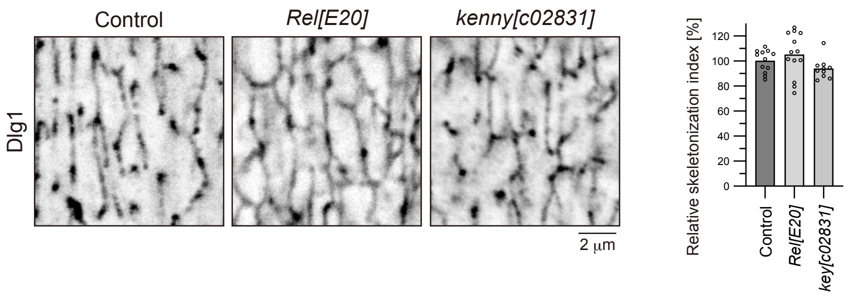


**Figure S2. T-tubule formation is independent of the NF-κB signaling pathway**
Images showing anti-Dlg1 staining of 3IL BWMs in the indicated genotypes with the relative skeletonization index quantified.

### Figure S2

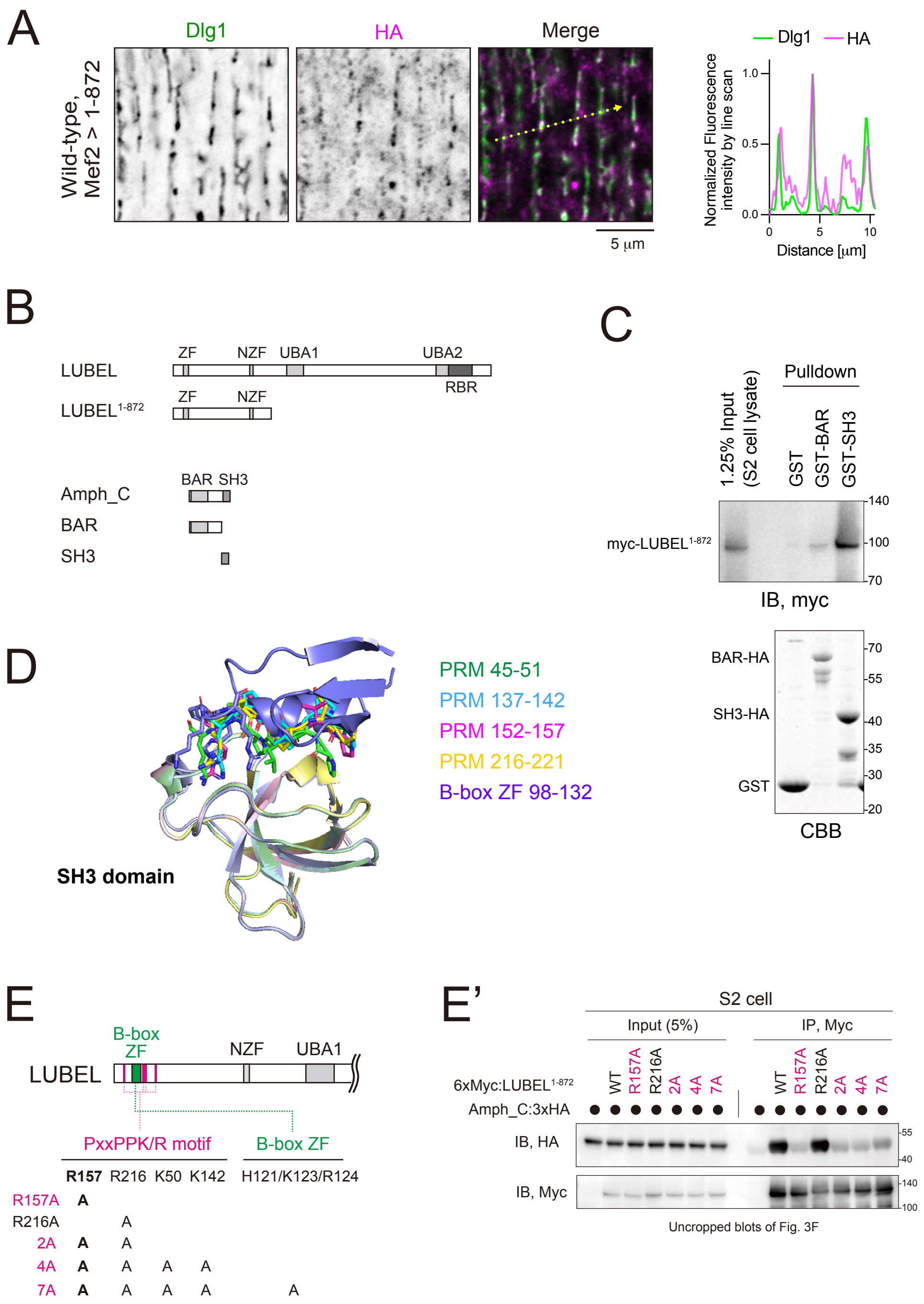


**Figure S3. Characterization of the interaction between LUBEL and Amph**
(A) Localization of HA-LUBEL1-872. 3IL BWMs expressing HA-LUBEL1-872 were stained with anti-Dlg1 and anti-HA antibodies. (B) Schematic representation of LUBEL and Amph truncations. (C) GST pulldown assay of truncated Amph and LUBEL constructs. Myc-LUBEL1-872 was expressed in S2 cells. Lysates containing Myc-LUBEL1-872 were incubated with beads conjugated to GST-fused truncated Amph isoform C (GST-BAR and GST-SH3). The resultant samples were immunoblotted with anti-Myc antibody or stained with CBB. (D) Structure of the Amph SH3 and LUBEL-N fragment complex using AlphaFold 2. Four proline-rich motifs and a B-box zinc finger are shown. (E) Schematic representation of LUBEL point mutants. (E') Co-IP assay of the LUBEL point mutants and Amph. The Myc-LUBEL1-872 fragments harboring the point mutations indicated in (E) were co-expressed with Amph isoform C-HA in S2 cells. Lysates were subjected to anti-Myc IP and immunoblotted for anti-Myc and anti-HA antibodies.

### Figure S3

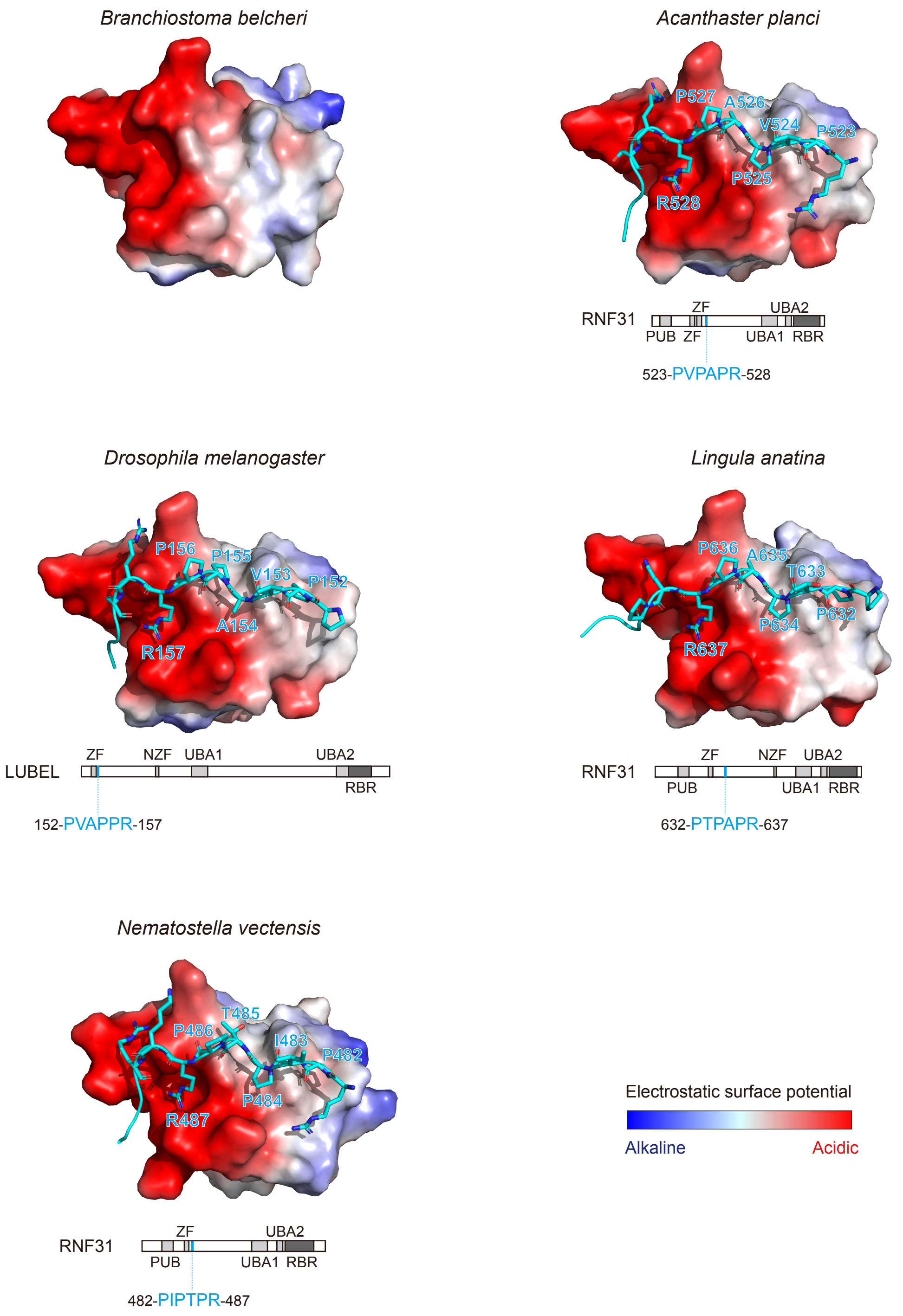


**Figure S4. AlphaFold prediction of RNF31-Amph interaction**
Predicted structures of the Amph SH3–RNF31 complex for each species shown in Figure 5F, generated using AlphaFold3. The Amph SH3 domain is depicted with its surface electrostatic potential, while RNF31/LUBEL is represented as cyan stick models. Below each structure, the domain organization of RNF31/LUBEL from each species is shown, with the Amph-interacting region (proline-rich motif) highlighted in cyan, along with its position and sequence.

### Figure S4

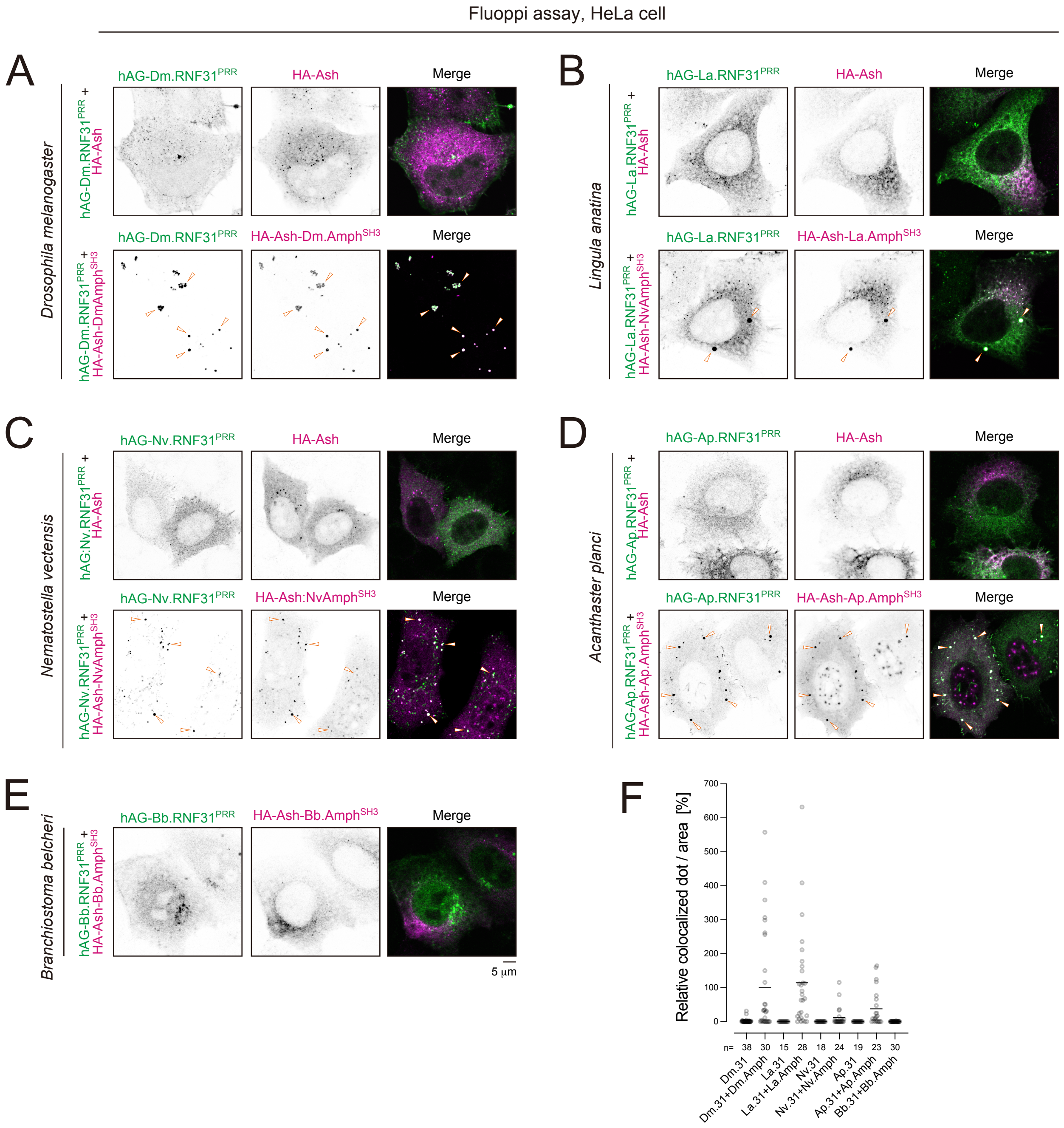


**Figure S5. Protein-protein interaction assay using Fluoppi system**
(A-E) Fluoppi analysis of the interaction between RNF31 and Amph in HeLa cells. HeLa cells transiently expressing the indicated constructs from each species listed were stained with anti-HA antibodies. (F) Quantification of foci positive for both RNF31 and Amph.

### Figure S5

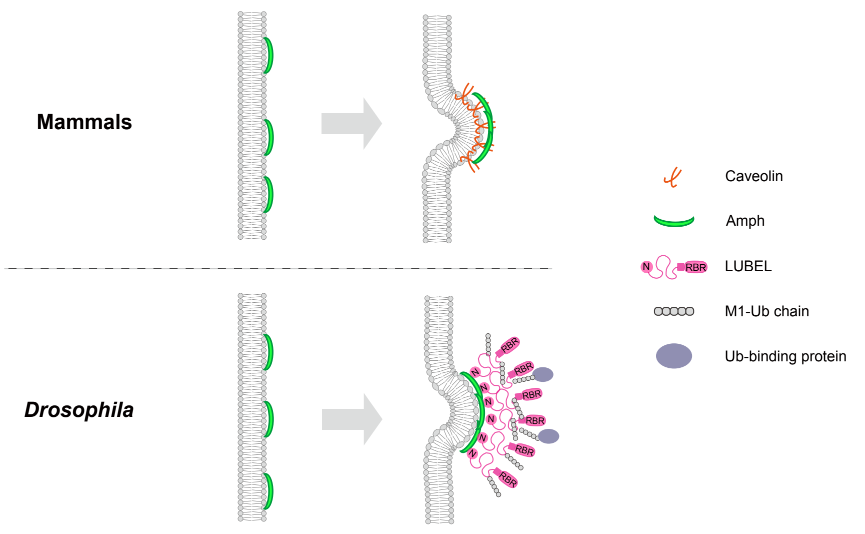


**Figure S6. Schematic models of the early stages of T-tubule biogenesis in mammals and *Drosophila***

### Figure S6
